# Progesterone is an Inducement of Heritable Pulmonary Arterial Hypertension with BMPR2 Mutation

**DOI:** 10.1101/2023.04.21.537897

**Authors:** Wei-Ping Hu, Si-Jin Zhang, Yong-Jie Ding, Jie Fang, Ling Zhou, Si-Min Xie, Xiao Ge, Li-Jun Fu, Qing-Yun Li, Jie-Ming Qu, Shan-Qun Li, Dong Liu

**Author notes:** Correspondence: Dong Liu, ^1^Department of Pulmonary and Critical Care Medicine, Ruijin Hospital, Shanghai Jiao Tong University School of Medicine, Shanghai, China, 200025 ^2^Institute of Respiratory Diseases, Shanghai Jiao Tong University School of Medicine, Shanghai, China, 200025, ^3^Shanghai Key Laboratory of Emergency Prevention, Diagnosis and Treatment of Respiratory Infectious Diseases, Shanghai, China, 200025,; Shan-Qun Li, Department of Pulmonary and Critical Care Medicine, Zhongshan Hospital, Shanghai Medical College, Fudan University, Shanghai, China, 200032.

## Abstract

**Background:** Bone morphogenetic protein type II receptor (BMPR2) gene mutation accounts for 80% of patients with heritable pulmonary artery hypertension (HPAH), and female mutation carriers have significantly higher penetrance rate than males. The inducement of HPAH penetrance and the mechanism of sex differential penetrance are still elusive.

**Methods:** We infected or transfected pulmonary artery smooth muscle cells (PASMCs) with shBMPR2 lentivirus or siBMPR2 to simulate the pathologic condition of BMPR2 heterozygous mutation and treated them with progesterone. The HPAH patient-derived induced pluripotent stem cells (iPSCs) were induced into vascular smooth muscle cells (VSMCs) to further verify the cellular phenotype. Wild-type flox^+/-^ female mice and SM22-cre BMPR2 flox^+/-^ female mice (CKO mice) were administered with 1-month progesterone, and their phenotype of PAH was evaluated by right heart catheterization and histopathological examination.

**Results:** Progesterone promoted the proliferation of PASMCs with BMPR2-knockdownby activating ERK pathway via progesterone receptor (PGR). Activated ERK not only upregulated the phosphorylation and elevation of cMYC, but also induced the transcription of endothelin (EDN1) by promoting the nuclear entry of c-JUN and combination on the its promoter region. Similar results were confirmed by iPSCs-VSMCs experiment. CKO mice developed PAH spontaneously and had increased expression of EDN1, which was further aggravated by exogenous progesterone.

**Conclusions:** Progesterone might be an inducement of HPAH penetrance caused by BMPR2 mutation, accounting for sex differential penetrance.

## Introduction

Pulmonary arterial hypertension (PAH) is a malignant cardiopulmonary vascular disease characterized by abnormal proliferation of pulmonary artery smooth muscle cells (PASMCs), with the stepwise development history of pulmonary vascular remodeling, elevated pulmonary arterial pressure, right heart failure and eventual death.^1^ Even with standardized drug therapy, the 5-year survival rate of adult PAH patients with medium risk is only 40-50%, and that of high-risk patients is only 10–20%.^2^

As a subtype of PAH, heritable PAH (HPAH) is an autosomal dominant disease with incomplete penetrance. The heterozygous mutation in the Bone morphogenetic protein type II receptor (BMPR2) gene is the main genetic pathogenic factor of PAH, accounting for 80% of patients with HPAH and 20% of patients with idiopathic PAH (IPAH).^3, 4^ BMPR2 is pivotal in maintaining pulmonary vascular homeostasis. Bone morphogenetic proteins (BMPs) induce the formation of tetramers of BMPR2 and type I receptors and induce the phosphorylation and nuclear translocation of smad1/5/8, thereby promoting the transcription of target genes and inhibiting cellular proliferation.^5^ The penetrance rate of male BMPR2 mutation carriers was approximately 14%, while that of female carriers was as high as 42%.^6^ This difference implies that female mutation carriers have a higher risk of developing PAH in life than male mutation carriers. However, the inducement of HPAH penetrance caused by BMPR2 mutation is not fully understood, and the mechanism of sex differential penetrance is also elusive.

In the clinical setting, the median age of PAH in women is 36 years old ^7^ and many female mutation carriers develop HPAH after gestation.^8^ Pregnancy is the most remarkable period for the fluctuation of sex hormones for women. The levels of 17β– estradiol and progesterone incrementally increase during gestation, peaking at more than 100 times that of non-pregnant period.^9^ In addition, the effect of progesterone in PAH is also unclear, and its relationship with BMPR2 in pulmonary vascular cells was not reported. Limited studies showed that progesterone could dilate pulmonary vasculature, and prophylactic administration of progesterone could alleviate the severity of PAH in rats after ovariectomy.^10, 11^ A clinical study reported that the low level of progesterone was associated with high mortality in women of reproductive age with IPAH.^12^ In summary, we speculated that progesterone might also have different effects on PASMCs in different BMPR2 mutation states and influence the occurrence and development of HPAH, besides from 17β-estradiol. We infected or transfected PASMCs with shBMPR2 lentivirus or siBMPR2 to simulate the pathologic condition of BMPR2 heterozygous mutation and explored the influence and mechanism of progesterone on them *in vitro*. The HPAH patient-derived induced pluripotent stem cells (iPSCs) were induced into vascular smooth muscle cells (VSMCs) to further verify the cellular phenotype. Wild-type flox^+/-^ mice and SM22– cre BMPR2 flox^+/-^ mice were bred and administered with progesterone to explore the effects of progesterone on pulmonary artery remodeling and right heart pressure *in vivo*.

## Methods

### Enzyme-linked immunosorbent assay (ELISA)

Human BMPR2 ELISA Kit (orb406355, Biorbyt, United Kingdom) was used to detect the level of soluble BMPR2 in serum. Its detection range was 6.25 pg/ml – 400 pg/ml, with the sensitivity of 1.56 pg/ml. To reduce the interference of albumin, the serum was diluted by 1.2 times with the dilution buffer. Experiment procedure and calculation method were referred to the instruction.

### Reagent and cell culture

Progesterone (S30586, Yuanye Bio, Shanghai, China) was dissolved in methanol at the stocking concentration of 10mg/ml. We used the final concentration of 100nM and 1μM, according to the highest level of progesterone in the third trimester and the other study.^14^ Similarly, β-estradiol (E2758, sigma, Germany) was dissolved in ethanol at the stocking concentration of 1mg/ml. The final concentration for RNA-seq was set as 100nM according to other studies.^15, 16^ PD0325901 (M1763, Abmole, USA) was the inhibitor of MEK pathway and was dissolved in dimethyl sulfoxide (DMSO, D2650, sigma, Germany). Bosentan (HY-A0013, MCE, China) was the inhibitor of endothelin receptor A/B and was dissolved in DMSO. Mifepristone (M3510, Abmole, USA) was the inhibitor if progesterone receptor and was dissolved in DMSO.

The protocol of culturing human PASMCs (3110, Sciencell, USA) was reported in our previous study. ^17^ To generate induced pluripotent stem cells (iPSCs), peripheral blood mononuclear cells (PBMCs) were harvested from 1 HPAH patient with BMPR2 exon1 189-196 deletion and 1 healthy subject. The establishment of healthy and patient-specific iPSC lines was performed in a GMP facility of Help Stem Cell Innovations Co., Ltd. (Nanjing, China) and the protocol was reported in our previous study.^18^ Several pluripotent markers (SSEA4, Oct4 and Nanog) were identified by flow cytometry and immunofluorescence stain (IF). The iPSCs were cultured in Stemflex Medium Combo Kit (15040066, Thermofisher, USA) and the culture protocol was followed according to its instructions. The directed differentiation of iPSCs towards vascular smooth muscle cells (VSMCs) was induced according to the authoritative paper and identified by IfF of αSMA. ^19^

### Lentivirus infection and transfection

The shRNAs of BMPR2 and lentivirus were provided by GeneChem Co., Ltd. (Shanghai, China). Preliminary experiments demonstrated that the optimal multiplicity of infection (MOI) of PASMCs was 30. PASMCs were seeded in the 6– well plate at the density of 30%, and then infected with lentivirus. After 12 hrs, we changed the medium and extracted RNA and protein at 72 hrs. Because shBMPR2-3 had the best efficacy of BMPR2 knockdown among the 3 shRNAs, it was used for RNA sequencing. (Data not shown)

According to the manufacturer’s instructions, siNC (50 nM), siBMPR2 (50 nM) and the siPGR (200 nM) were transfected into PASMCs by Opti-MEM (31985088, Thermofisher, USA) and Lipofectamine 3000 (L3000015, Invitrogen, UK). The siRNAs were synthesized by RiboBio Co., Ltd. (Guangzhou, China), and their sequences were listed in the **Supplementary Table 1**.

### RNA sequencing and bioinformatic analysis

Flowchart of sample treatment and collection was shown in the Supplementary Figure 1c. Each treatment had three replicate wells. RNA sequencing was performed in GeneChem Co., Ltd. (Shanghai, China). A total amount of 1 µg RNA per sample was used as input material for the RNA sample preparations. Sequencing libraries were generated using NEBNext® UltraTM RNA Library Prep Kit for Illumina® (NEB, USA) following manufacturer’s recommendations. The clustering of the index-coded samples was performed on a cBot Cluster Generation System using TruSeq PE Cluster Kit v3-cBot-HS (Illumia) according to the manufacturer’s instructions. After cluster generation, the library preparations were sequenced on an Illumina Novaseq platform and 150 bp paired-end reads were generated.

FPKM, expected number of Fragments Per Kilobase of transcript sequence per Millions base pairs sequenced, is used for quantification of gene expression level. FeatureCounts v1.5.0-p3 was used to count the reads numbers mapped to each gene. And then FPKM of each gene was calculated based on the length of the gene and reads count mapped to this gene.

Gene Ontology (GO) enrichment analysis of differentially expressed genes was implemented by the cluster Profiler R package, in which gene length bias was corrected. GO terms with corrected *P* value less than 0.05 were considered significantly enriched by differential expressed genes. KEGG is a database resource for understanding high-level functions and utilities of the biological system. (http://www.genome.jp/kegg/) We used cluster Profiler R package to test the statistical enrichment of differential expression genes in KEGG pathways. Venn graph was made by a website tool of bioformatics analysis. (https://www.xiantao.love/)

### Cell viability and migration assay

After transfected with siRNA for 36 hrs, PASMCs were incubated with progesterone (100nM & 1μM) and other drugs for functional experiments. Alarmar blue assay (40202ES60, Yeasan, Shanghai, China) and EdU assay (CX002, EpiZyme, Shanghai, China) were performed to evaluate the ability of cell proliferation. In addition, the ability of cell migration was detected by Transwell assay (TCS003024, Jet BIOFIL, Guangzhou, China). Alarmar blue asaay had at least four duplicates, and EdU assay and Transwell assay had three duplicates per group. Detailed experiment procedures were reported in our previous study.^17^

### Quantitative polymerase chain reaction (qPCR)

The remained samples of RNA sequencing were used for reverse transcription and qPCR to validate the sequencing result. After transfected with siRNA for 48 hrs in 12–well plates, PASMCs were incubated with progesterone (1μM), PD0325901 (10μM) or Mifepristone (100nM) for 2 hr, and then extracted for RNA. The qPCR experiment had three duplicates per group. Reagent information was reported in our previous study ^17^ and primer sequence was listed in the **Supplementary Table 2.**

### Western blot

PASMCs were transfected with siRNA for 48 hrs in 6-well plates, then incubated with progesterone (1μM) or other drugs for 2 hr or 24hr, and finally harvested for protein extraction. Lung tissues of solvent or progesterone-treated transgenic mice were also harvested by lysis buffer and minced by Tissue Lyser® and Ultrasonic Processor. Detailed information of reagent and instrument was reported in our previous studies ^17, 20^ and information of antibodies was listed in the **Supplementary Table 3**.

### Immunofluorescence stain (IF)

Transfected PASMCs were seeded in Millicell® EZ SLIDES (C86024, Merck, Germany) at the density of 8 × 10^3^ cells, and stimulated by progesterone (1μM) for 2h. The iPSCs were labeled for pluripotentmarker Oct4 and nucleiwere labeled with Hoechst stain (Blue). We used the immunofluorescence application solutions kit (12727, CST, USA) and followed its instruction. Paraffin sections of lung tissues were co-stained with vWF and αSMA using the IF method.

### Flow Cytometry

The iPSCs were digested by accutase, gently washed with phosphate-buffered saline and then labeled for pluripotentmarkers SSEA4, Oct4 and Nanog, respectively. We reported the information of antibodies in the **Supplementary Table 3**. Single-cell suspensions of iPSCs were streamed in the BD FACSAria™II flow cytometer (BD bioscience Inc., USA), and FlowJo 10 software (BD bioscience Inc., USA) was used to to evaluate the percentage of positive cells.

### Chromatin immunoprecipitation (ChIP)

We used the UCSC Genome Browser (http://www.genome.ucsc.edu/) and its embedded JASPAR database to predict whether AP-1 could combine on the promoter region of ECE1 and EDN1. PASMCs were plated into 10 cm culture dishes to achieve the density of 90%, and then collected. SimpleChIP® Plus Enzymatic Chromatin IP Kit (Magnetic Beads) (9005, CST) was used for ChIP experiment based on its instruction. Because IF results showed that c-Jun was mainly distributed in nucleus, we used c-Jun rabbit mAb (9165, 1:50, cst) as the bait antibody and normal rabbit IgG (2729, cst) as the negative control. Information of primers was listed in the **Supplementary Table 2.**

### DNA extraction, sanger sequencing and agarose gel electrophoresis

Genomic DNA extraction kit (DP304, TIANGEN, China) was used to collect the DNA of iPSCs and mouse tails according to its instruction. We used 2×Hieff Canace® Plus PCR Master Mix (10154ES08, YEASAN, China) for polymerase chain reaction (PCR), following its instruction. Sanger sequencing was performed in Tsingke Co., Ltd. (Beijing, China) to examine the BMPR2 exon1 mutation region of HPAH-iPSCs. Agarose gel electrophoresis was performed to identify the genotypes of neonatal mice. The primer sequences of BMPR2 exon1, BMPR2 flox and SM22-Cre were listed in **Supplementary Table 2.**

### Animal breeding, identification and intervention

BMPR2 flox ^+/-^ mice were gifted from the lab of Prof. Qi-qun Tang, and crossed with SM22-Cre mice to acquire the SM22-Cre BMPR2 flox ^+/-^ mice.^21^ SM22-Cre mice (C001004) were provided by Cyagen Co., Ltd. (Suzhou, China). The mice were routinely bred in the SPF (Specified Pathogen Free) facility of Cyagen Co., Ltd. (Suzhou, China) and genetically identified at two weeks of age. At four months of age, they were transferred to the SPF facility of Charles River Co., Ltd. (Shanghai, China) for drug administration. Progesterone was solved in corn oil (C8267, sigma, Germany) at the concentration of 20mg/ml with the aid of ultrasound. The weight of female mice at four months of age was uniformly regarded as 30g. After adapt to the new environment for 1 week, 100μl of progesterone solution (60mg/kg) or corn oil were subcutaneously injected to the neck of mice per week for four times.^22^ To confirm the efficacy, progesterone solution was freshly prepared every week. Weighing and phenotype detection were taken one week after the last drug administration.

### Echocardiography and Right heart catheterization (RHC)

After shaving the hair on the chest in advance, mice were induced anesthesia by inhaling 3% isoflurane and then maintained by 1–1.5% isoflurane to keep the heart rate around 400 beats/min. After anesthesia, they were gently fixed on a thermostatic plate in a supine position. All the echocardiograms were obtained by a skilled and blinded animal technician using a 40MHz probe (Vevo®2100, VisualSonics Co., USA). We measured right ventricular wall thickness at the right parasternal long-axis view.

Mice were anesthetized using inhaled isoflurane at 3% concentration. To measure right ventricular systolic pressure (RVSP), we used the Millar Mikro-Tip® single pressure catheter (SPR-671, Millar, USA) to enter the right ventricle through the right external jugular vein. The stable waveform of RVSP was recorded by the pressure transducer at the anterior end of catheter, transformed and calculated by the BL-420S Bio Lab System (Chengdu TME Technology Co., Ltd., Chengdu, China).

### Hematoxylin–eosin (H&E) stain and Immunohistochemical stain (IHC)

After the RHC experiment, the mice were euthanized by intraperitoneally injecting excessive sodium pentobarbital (10011014, YUYAN INSTRUMENTS Co., China). The lung and heart of each mouse were collected after flushing blood. The right lung was frozen by dry ice and stored at −80°C for qPCR and Western blot. The left lung and a part of hearts were fixed by 4% paraformaldehyde at room temperature for 24 hr and embedded in paraffin. The weight ratio of the RV wall to the left ventricle plus septum (RV/LV+S) was measured to evaluate RV hypertrophy, and then they were frozen by dry ice. Detailed protocols of H&E staining and IHC were reported in our previous study, ^23^ and the list of antibodies was shown in the **Supplementary Table 3.** CaseViewer software (3DHISTECH Ltd., Hungary) was used to measure the indices of H&E stain, including mean wall thickness of small arteries (diameter < 50μM or 50-100μM), RV wall area and RV cavity area. Image J software (NIH, Bethesda, USA) was used to analyze the average optical density of BMRP2 in pulmonary arterioles.

### Statistical analysis

As for the clinical research of soluble BMPR2, the continuous variables were expressed as the median and interquartile range (IQR). The categorical variables were expressed as number (percentage), and analyzed by Chi-square test. As for cellular and animal experiments, all the data were expressed as the mean ± standard deviation (SD) from at least three replicates. We used the Mann–Whitney test or for two groups’ comparison and Kruskal–Wallis test for multiple groups’ comparison with Dunnett test for pairwise comparisons. The specific number of animals of each group was reported in the figure legend. IBM SPSS statistics 23 (SPSS Inc., Chicago, IL, USA) and GraphPad Prism 6 (GraphPad Software, CA, USA) were used to perform the above statistical analysis. The receiver operating characteristic (ROC) curve was performed by MedCalc software v19.5.3 (MedCalc Software Ltd, Ostend, Belgium). A two-tailed *P* < 0.05 was regarded as statistical significance.

## Result

### Progesterone promoted the proliferation of BMPR2-knockdown PASMCs

The strategy for RNA sequencing was shown in **Figure 1a**. PASMCs were infected with lentiviruses containing the shBMPR2-GFP plasmid. After 48 hours of infection, normal and BMPR2-knockdown PASMCs were stimulated with 17β-estradiol or progesterone for 24 hours. Gene Ontology (GO) enrichment analysis of differentially expressed genes (DEGs) showed that 17β-estradiol and progesterone increased the ranking of proliferation and migration gene sets in the BMPR2-knockdown PASMCs significantly. (**Figure 1b & Supplementary Figure 1a**) Compared with 17β-estradiol, progesterone had an obvious effect on the gene set of proliferation based on the rank. In addition, heatmap and principal component analysis of DEGs demonstrated that 17β-estradiol and progesterone had similar effects on gene transcription in PASMCs. However, the effects of sex hormones on gene transcription in PASMCs were distinct between normal and BMPR2-knockdown conditions, suggesting that sex hormones might have additional pathogenic mechanisms in PAH patients with BMRP2 mutations. (**Supplementary Figure 1b-c**)

**Figure 1.**
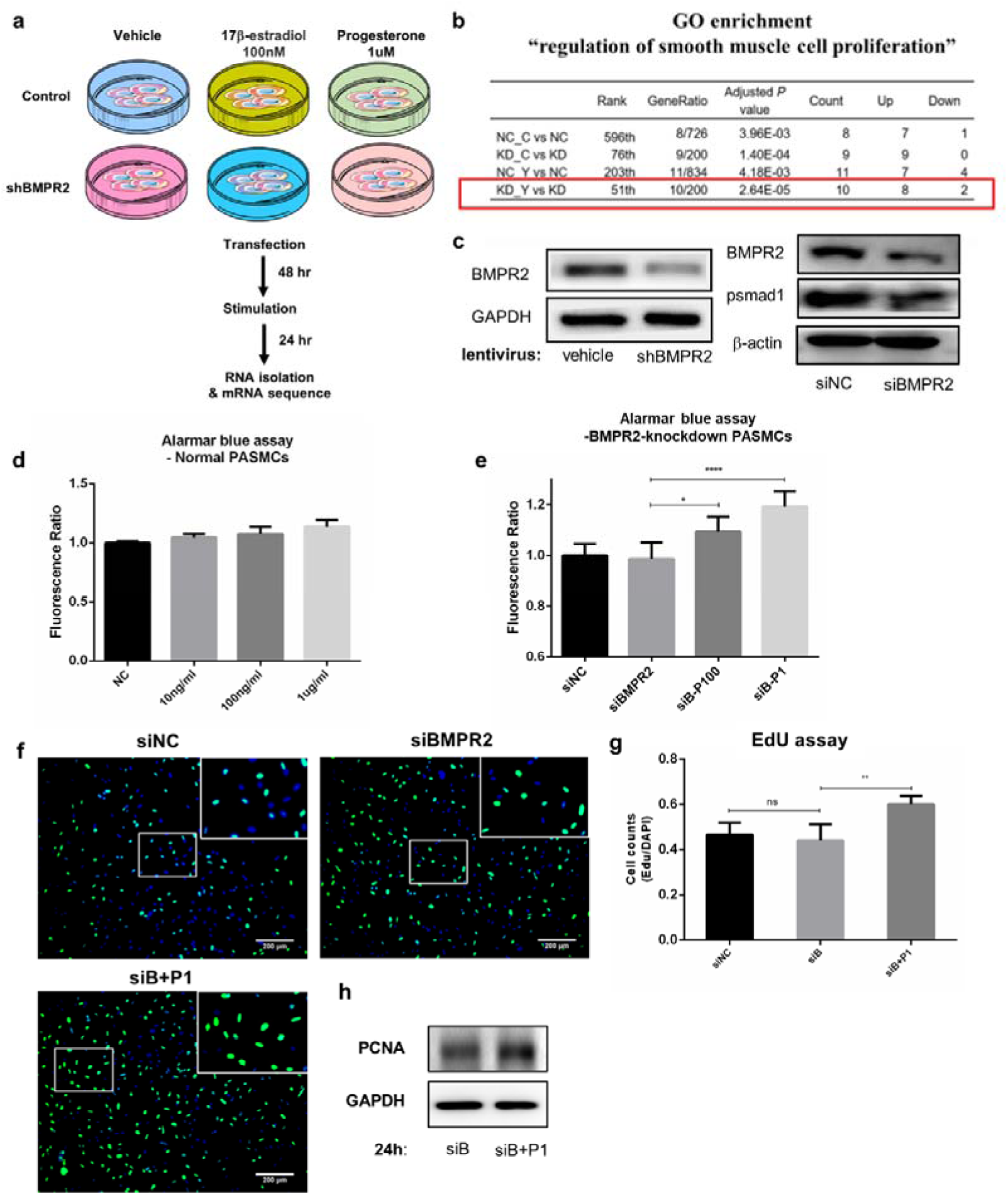
Progesterone promoted the proliferation of BMPR2-knockdown PASMCs but not normal PASMCs. (**a**) Schematic diagram of sample processing for RNA sequencing. (**b**) Under the condition of BMRP2 knockdown, the stimulation of progesterone promoted the enrichment of pro-proliferative genes in PASMCs. (**c**) Both infecting shBMPR2 lentivirus (MOI=30, 48 hr) and transfecting siBMPR2 (50 nM, 48 hr) effectively decreased the protein level of BMPR2 in PASMCs. (**d**) Progesterone had little effect on the proliferation of normal PASMCs. (**e-h**) Under the condition of BMPR2 knockdown, progesterone (for 24 hr) significantly promoted the proliferation of PASMCs. (e) used Alarmar blue assay, (f-g) were EdU assay and its statistical graph, and (h) was the expression of PCNA. **Abbreviation**: PASMCs, pulmonary artery smooth muscle cells; NC, normal PASMCs; NC_C, normal PASMCs stimulated with 17β-estradiol; NC_Y, normal PASMCs stimulated with progesterone; KD, shBMRP2 lentivirus-infected PASMCs; P100, 100nM of progesterone; P1, 1μM of progesterone; PCNA, proliferating cell nuclear antigen. ns, non-significance; *, *P*<0.05; **, *P*<0.01; ****, *P*<0.001.

To downregulate BMPR2 mRNA transiently, we transfected PASMCs with siRNA and collected proteins after 72 hr for Western blot. **Figure 1c** showed that BMPR2 was successfully downregulated by shBMPR2-lentiviruses and siBMPR2, respectively. In normal and siBMPR2-transfected PASMCs (48 hr), we incubated them with 17β-estradiol or progesterone (24 hr), and detected the proliferative ability using Alarmar blue assay. Both17β-estradiol and progesterone had little pro-proliferative effect on normal PASMCs. (**Supplementary Figure 2a and Figure 1d).** As for BMPR2-knockdown PASMCs, progesterone (100nM and 1μM) could promote their proliferation (**Figure 1e**), but 17β-estradiol without any significant effect (**Supplementary Figure 2b**). Western blot of PCNA and EdU staining also proved that 1μM of progesterone could promote the proliferation of BMPR2-knockdown PASMCs. (**Figure 1f–h**) Transwell assay demonstrated that progesterone (100nM and 1μM) could promote the migration of BMPR2-knockdown PASMCs. (**Supplementary Figure 2c-d**)

### Progesterone promoted the proliferation of BMPR2-knockdown PASMCs via the PGR-ERK-cMYC/EDN1 axis

To explore the mechanism by which progesterone promoted the proliferation of BMPR2-knockdown PASMCs, we performed KEGG enrichment analysis of DEGs. (**Figure 2a**) By comparing two groups of “BMRP2 knockdown + progesterone” and “BMRP2 knockdown” (“KD_Y vs KD”), we found that the MAPK pathway was significantly enriched. Western blot confirmed that 1μM of progesterone could up-regulate pERK and pJNK at 2 hr and 6 hr in siBMPR2-transfected PASMCs. (**Figure 2b**) In comparison with the JNK pathway, the ERK pathway was more closely related to cell proliferation, so we mainly investigated the ERK pathway in the subsequent study.^24^

**Figure 2.**
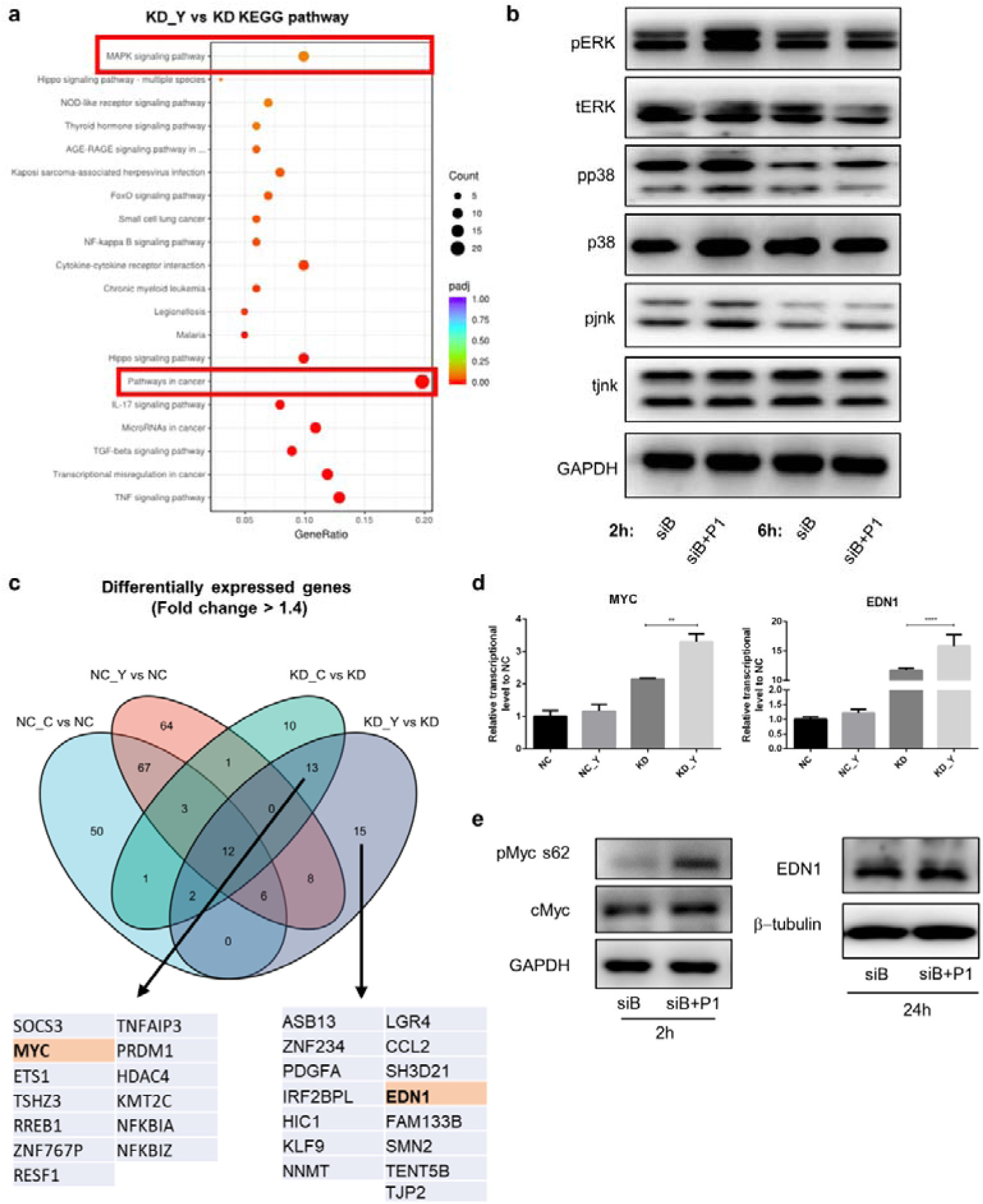
Progesterone promoted phosphorylation of ERK and upregulated cMYC and EDN1 in BMPR2-knockdown PASMCs. (**a**) Progesterone (24 hr) enriched the genes of “MAPK pathways” and “Pathways in cancer” in BMRP2-knockdown PASMCs. (**b**) Progesterone (for 2 hr) activated the ERK pathway in BMPR2-knockdown PASMCs. (**c**) Venn graph, which brought into all the differentially-expressed genes with the fold change > 1.4, showed that MYC and EDN1 were upregulated by progesterone (24 hr) only under the condition of BMPR2 knockdown. (**d-e**) Progesterone upregulated the mRNA and protein of c-MYC and EDN1 in BMPR2-knockdown PASMCs. **Abbreviation**: ERK, extracellular regulated protein kinases; EDN1, endothelin1; MAPK, mitogen-activated protein kinase. **, *P*<0.01; ****, *P*<0.001.

Moreover, “Pathways in cancer” was the most enriched pathway, in which MYC was one of the enriched genes. (**Figure 2a**) MYC is a well-known proto-oncogene, and its mutation is related to carcinogenesis of various tissue cells, which might be the possible mechanism by which progesterone promoted proliferation.^25^ Similarly, the Venn graph also showed that MYC was obviously up-regulated in the comparison of “KD_Y vs KD”, but it was not significantly changed by sex hormones in the normal PASMCs. (**Figure 2c**) In addition, we found that EDN1 (encoding Endothelin-1) was only significantly increased in n the comparison of “KD_Y vs KD”. (**Figure 2c**) As a strong vasoconstrictor, Endothelin-1 (EDN1) plays an important pathogenic role in PAH. Endothelin receptor inhibitors (such as Bosentan) are one of the classic targeted drugs for PAH.^26^ Therefore, we regarded EDN1 as a potential mechanism by which progesterone promoted the proliferation of BMPR2-knockdown PASMCs. We performed cell experiments to verify the results of RNA sequencing. (**Figure 2d–e**) qPCR showed that progesterone increased the mRNA expression of MYC and EDN1 in shBMRP2-transfected PASMCs. (**Figure 2d**) Western blot showed progesterone promoted the phosphorylation of cMyc at serine 62 and the upregulation of cMyc and EDN1 at 2 hr and 24 hr, respectively. (**Figure 2e**) Correspondently, a previous study reported that phospho-Myc Ser62 could improve the stability of cMyc and inhibit its degradation.^27, 28^

To verify the pro-proliferative effect of activated ERK pathway and EDN1, BMPR2-knockdown PASMCs were simultaneously incubated with progesterone, PD0325901 or Bosentan in BMPR2-knockdown PASMCs for 24 hr. Alarmar blue assay (**Figure 3a**) and EdU assay (**Figure 3b–c**) were performed to evaluate the proliferation ability. 0.1% DMSO was the solvent control. Low concentrations of PD0325901 (1μM) and Bosentan (10μM & 20μM) could reverse the pro-proliferative effect of progesterone in BMPR2-knockdown PASMCs, and high concentrations of PD0325901 (10μM) and Bosentan (50μM) directly inhibited the proliferation.

**Figure 3.**
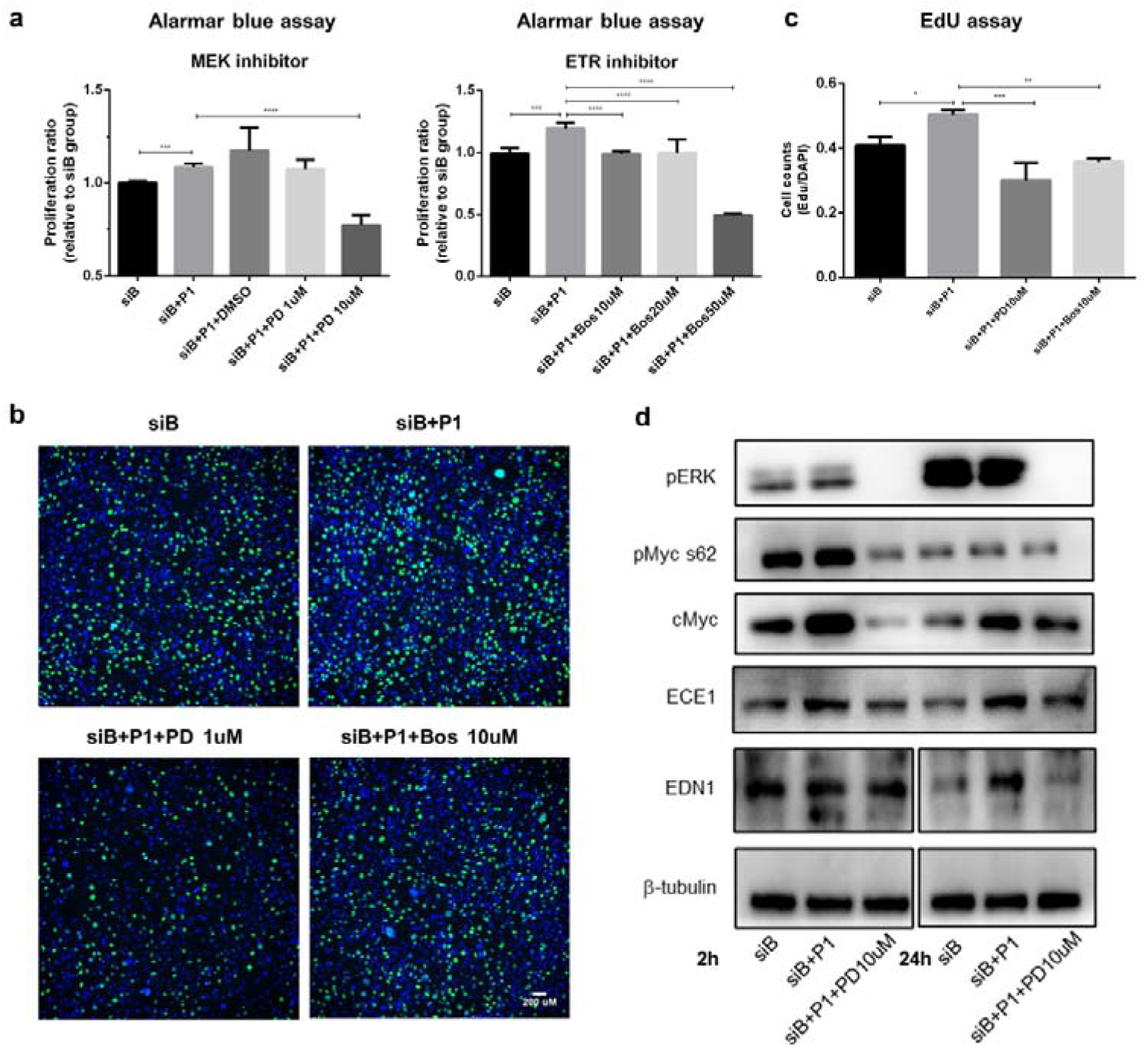
Progesterone promoted the proliferation of BMPR2-knockdown PASMCs via the ERK-cMYC/EDN1 axis. (**a**) Alarmar blue assay showed that PD0325901 and Bosentan (for 24 hr) reversed the pro-proliferative effects of progesterone in BMPR2-knockdown PASMCs**. (b-c**) EdU assay showed that PD0325901 and Bosentan (for 24 hr) also reversed the effects of progesterone. (**d**) Inhibiting ERK pathway by PD0325901 reversed the upregulation of these proteins induced by progesterone in BMPR2-knockdown PASMCs. **Abbreviation**: ECE1, endothelin converting enzyme 1; PD, PD0325901; Bos, Bosentan. ns, non-significance; *, *P*<0.05; **, *P*<0.01; ***, *P*<0.001; ****, *P*<0.001.

To explore the regulatory relationship between the activation of ERK pathway and the increase of cMyc and EDN1, BMPR2-knockdown PASMCs were simultaneously incubated with progesterone and PD0325901 (the inhibitor of ERK pathway). ^27^ ECE1 (encoding endothelin converting enzyme 1) which could cleave the inactive short precursor big EDN1 into mature EDN1 ^29^ had also been detected in this experiment. The upregulation of pMyc Ser62, cMyc and ECE1 induced by progesterone at 2 hr was reversed by PD0325901. (**Figure 3d**) Similarly, the upregulation of cMyc, ECE1 and EDN1 induced by progesterone at 24 hr were also reversed by PD0325901. (**Figure 3d**)

To identify whether progesterone played a role through the progesterone receptor (PGR), we used siRNA to transiently downregulate PGR. Results of qPCR and Western blot showed that the expression of PGR successfully declined after transfection. (**Figure 4a&b**) Alarmar blue assay and EdU assay demonstrated that the pro-proliferative effect of progesterone on BMPR2-knockdown PASMCs was reversed by PGR knockdown, indicating that progesterone acted through PGR. (**Figure 4c–e**) In addition, **Figure 4f** showed that knockdown of PGR could also reverse the up-regulation effect of progesterone on pERK and EDN1, suggesting that PGR was located in upstream of ERK pathway.

**Figure 4.**
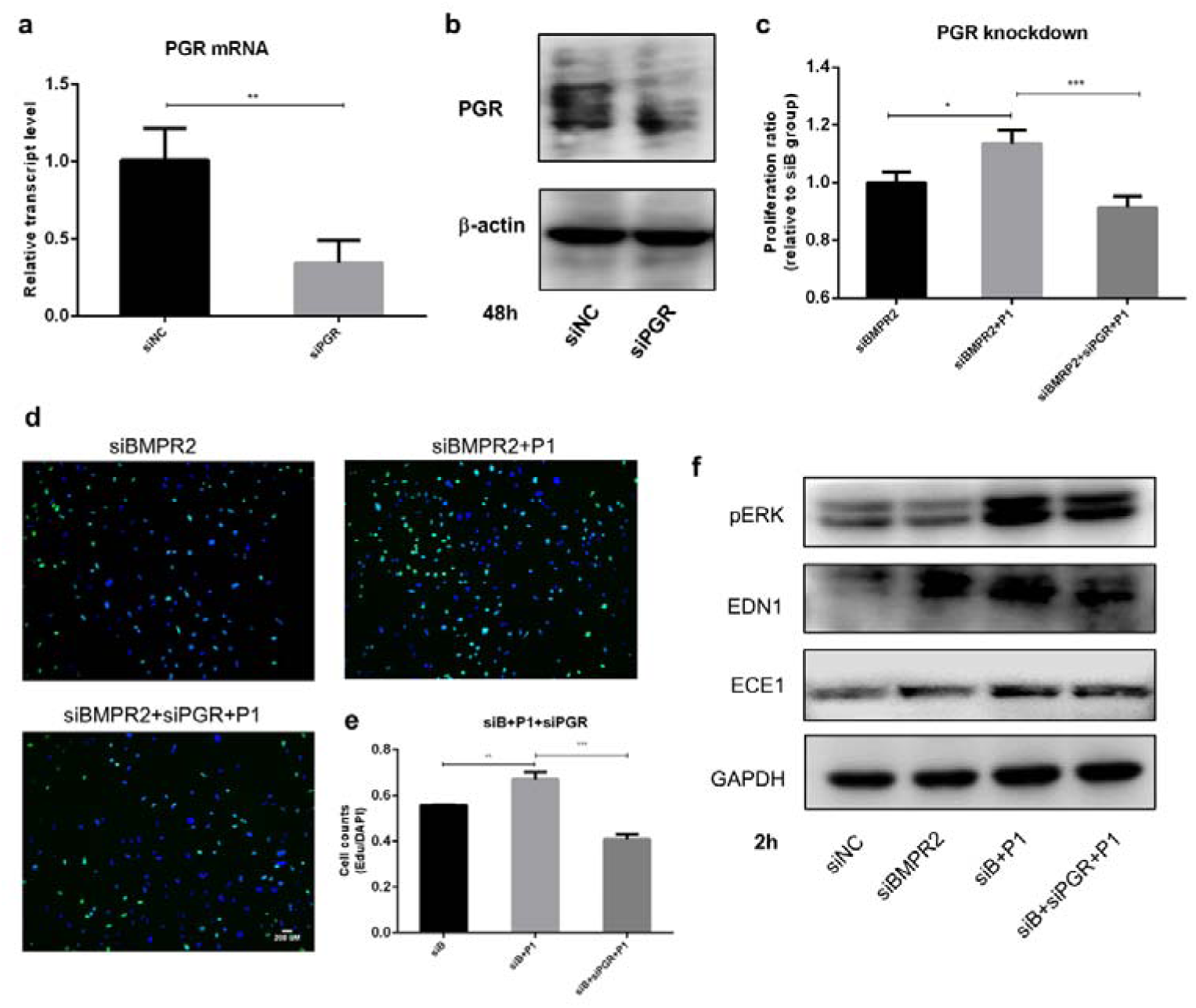
Progesterone promoted the proliferation of BMPR2-knockdown PASMCs dependent on PGR. (**a-b**) Transfecting siPGR (200nM, 72 hr) effectively downregulated the mRNA and protein of PGR. (**c-e**) Knockdown of PGR reversed the pro-proliferative effects of progesterone (for 24 hr) in BMPR2-knockdown PASMCs. (c) used Alarmar blue assay, and (d-e) were EdU assay and its statistical graph. (**f**) Knockdown of PGR reversed the upregulation of these proteins induced by progesterone (for 2 hr) in BMPR2-knockdown PASMCs. **Abbreviation:** PGR, progesterone receptor. *, *P*<0.05; **, *P*<0.01; ***, *P*<0.001.

### Progesterone induced the nucleus translocation of c-JUN and increased its combination on promoter region of EDN1

Activated ERK pathway promotes the translocation of c-JUN and c-FOS into the nucleus. The dimer of c-JUN and c-FOS is known as the transcription factor AP-1 which can bind onto the promoter region and promote transcription of targeted genes.^30^ To determine whether AP-1 was the key molecule of progesterone to promote proliferation, we detected the intracellular distribution of c-JUN and c-FOS, using Western blot of cytoplasmic and nuclear proteins and immunofluorescence. We found that c-JUN was mainly distributed in the nucleus of PASMCs, and c-FOS was mainly distributed in the cytoplasm of PASMCs. (**Figure 5a–c & Supplementary Figure 3a-b**) Western blot showed that progesterone promoted the nuclear entry of c-JUN and c-FOS, which was reversed by knockdown of PGR. (**Figure 5a**) By comparing the nucleus and cytoplasm ratio of mean fluorescence intensity (MFI) of c-JUN between different treatment groups, we demonstrated that progesterone upregulated the MFI of c-JUN in the nucleus, and knockdown of PGR could reverse this effect. (**Figure 5b&c**) As for c-FOS, we compared the percentage of cells with c-FOS distributed only in the nucleus, and found that progesterone increased the percentage of c-FOS into the nucleus, which was reversed by knockdown of PGR. (**Supplementary Figure 3a-b**)

**Figure 5.**
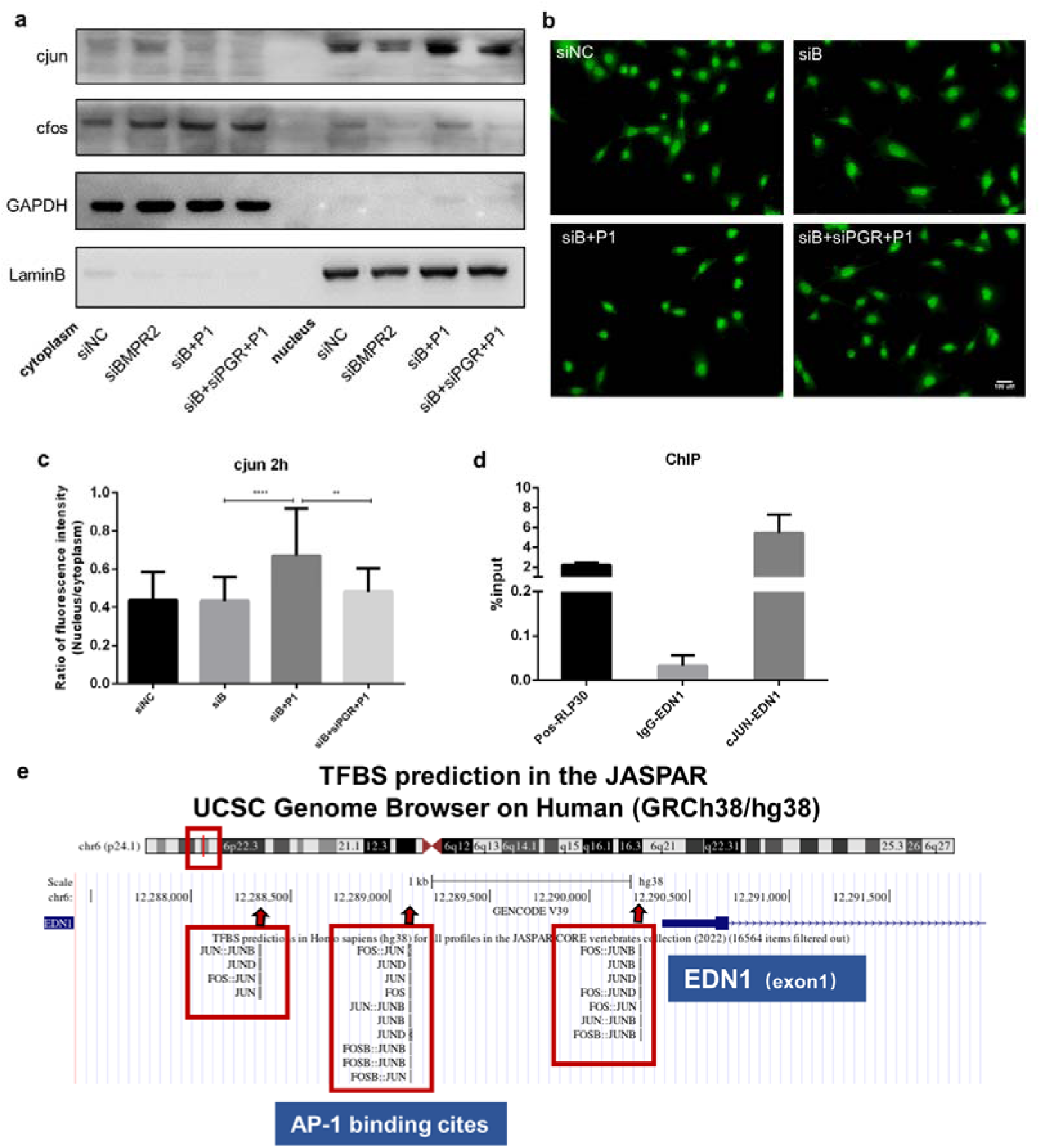
Progesterone promoted the entry of c-JUN into the nucleus which combined onto the promoter region of EDN1. (**a-b**) Progesterone (for 2 hr) promoted the nuclear translocation of c-JUN in BMPR2– knockdown PASMCs, which could be reversed by knockdown of PGR. (**c**) Nucleus and cytoplasm ratio of mean fluorescence intensity of c-JUN. (**d**) Compared with RLP30 (the positive control) and IgG (the negative control), the %input of cJUN was over 4%, demonstrating its binding to the promoter region of EDN1. (**e**) JASPAR database predicted that AP-1 (the dimer of c-JUN and c-FOS) could combine onto the promoter region of EDN1. **Abbreviation**: ChIP, chromatin immunoprecipitation; AP-1, activator protein-1. **, *P*<0.01; ***, *P*<0.001.

ChIP assay demonstrated that c-JUN could combine onto promoter region of EDN1. (**Figure 5d**) JASPAR database also predicted that AP-1 could combine onto the promoter region of ECE1 and EDN1. (**Figure 5e &Supplementary Figure 3c**) In summary, progesterone regulated EDN1 expression probably through PGR, promoted the entry of c-JUN into the nucleus, and acted by binding to the promoter region of EDN1.

Although we did not find the binding of c-JUN onto the ECE1 promoter region in the ChIP experiment (data not shown), but progesterone (2 hr) upregulated the mRNA level of ECE1 in BMPR2-knockdown PASMCs, which was reversed by PGR inhibitor (Mif) and ERK inhibitor (MEKi). (**Supplementary Figure 3d**) We speculated that progesterone might upregulate ECE1 through other mechanisms.

### ECE1 and EDN1 were evidently upregulated when BMPR2 was downregulated

As mentioned previously, progesterone did not influence the proliferation of normal PASMCs, but it promoted the proliferation of BMPR2-knockdown PASMCs (**Figure 2d–e**). Then, what is the intrinsic mechanism of declined BMPR2 that makes PASMCs more sensitive to the effect of progesterone? We transfected PASMCs with siNC and siBMPR2, extracted the protein after 48 hr, and performed Western blot. **Supplementary Figure 4a** showed that cMyc, ECE1, and EDN1 were increased in siBMRP2-transfected PASMCs significantly. We speculated that the inherently elevated expression of these key proteins might be responsible for the increased sensitivity of BMPR2-knockdown PASMCs to progesterone. In addition, we also performed qPCR on multiple batches of samples, and all of results repeated the up-regulation of ECE1 and EDN1 mRNA after BMRP2 knockdown. (**Supplementary Figure 4b**)

We also detected differences in the expression of tERK and their downstream transcription factors. (**Supplementary Figure 4c**) pERK (**Figure 4f**) and tERK (**Supplementary Figure 4c**) had no difference between two groups, suggesting ERK was not the reason for the difference in progesterone responses. c-JUN, c-FOS, and junB were not significantly up-regulated in siBMRP2-transfected PASMCs, so these transcription factors did not account for progesterone sensitivity. (**Supplementary Figure 4c**)

### Progesterone promoted the proliferation and migration of iPSCs-VSMCs derived from a PAH patient with BMPR2 mutation

To further confirm the above cellular phenotype and mechanism, we isolated PBMCs from a HPAH patient with heterozygous mutation of BMPR2 exon1 189-196 deletion (AAATGAAG) and reprogramed PBMCs into iPSCs. The iPSCs demonstrated high expression levels of pluripotent markers (SSEA4, Oct4 and Nanog). (**Supplementary Figure 5a&b**) We extracted DNA from the patient’s iPSCs and performed Sanger sequencing to confirm the remaining existence of BMPR2 mutation after induction. (**Supplementary Figure 5c&d**) Then, IPSCs were induced into VSMCs according to the flowchart shown in **Supplementary Figure 6a**. Different timepoints of micrographs showed obvious morphological differences during the period of induction. (**Supplementary Figure 6b**) Immunofluorescence of α-SMA on day 6 showed that almost all the cells were stained red, indicating that the induction efficiency was close to 100%. (**Supplementary Figure 6b**)

We induced iPSCs from a healthy person and the HPAH patient with BMPR2 mutation into VSMCs, respectively. After the stimulation of progesterone (1μM) for 24 hr or 8 hr, Alarmar blue assay or Transwell assay was performed. In normal iPSCs-VSMCs, progesterone had little effect on their proliferation and migration, but progesterone promoted the proliferation and migration of the HPAH patient’s VSMCs. (**Figure 6a–c**) Surprisingly, we did not observe the decrease of BMPR2 in the HPAH patient’s VSMCs in comparison with the normal VSMCs. (**Supplementary Figure 6c-e**) Moreover, progesterone did not significantly affect the expression of BMRP2 in both VSMCs, (**Supplementary Figure 6d**) suggesting the existence of other mechanism that made the HPAH patient’s VSMCs more sensitive to progesterone.

**Figure 6.**
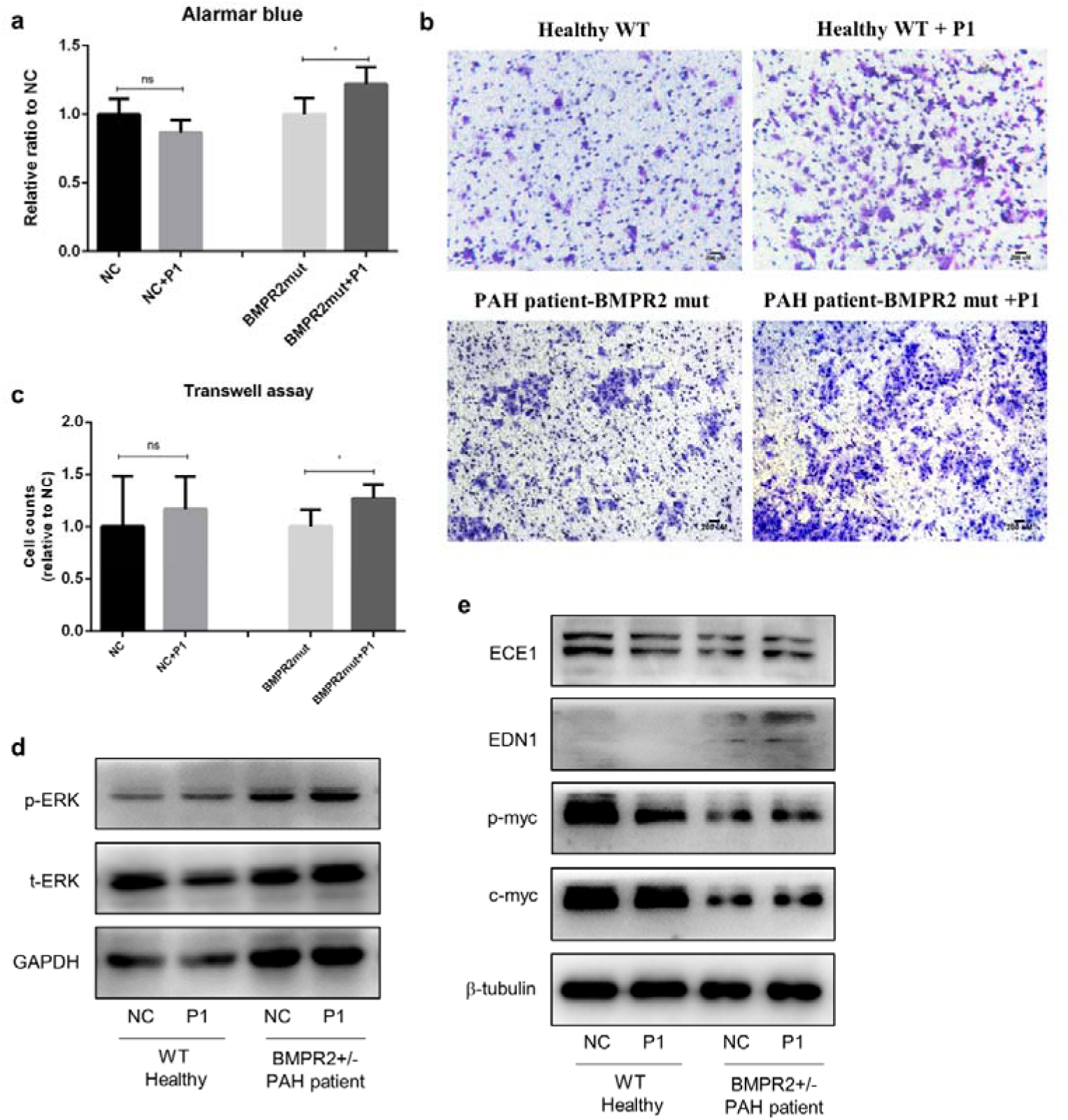
Progesterone promoted the proliferation and migration of iPSCs-VSMCs derived from a PAH patient with BMPR2 mutation. (**a**) Alarmar blue assay showed that progesterone (24 hr) upregulated the proliferation in iPSCs-VSMCs of a HPAH patient, but not influenced iPSCs-VSMCs of a normal person. (**b-c**) Transwell assay showed that progesterone upregulated the migration (8 hr) in iPSCs-VSMCs of a HPAH patient, but not influenced iPSCs-VSMCs of a normal person. (**d**) Progesterone (2 hr) induced the phosphorylation of ERK in iPSCs-VSMCs. (**e**) In comparison with normal iPSCs-VSMCs, EDN1 was significantly increased in PAH iPSCs-VSMCs that could be further upregulated by progesterone (24 hr). **Abbreviation:** iPSCs, induced pluripotent stem cells; VSMCs, vascular smooth muscle cells; HPAH, hereditary pulmonary artery hypertension; BMPR2 mut, BMRP2-mutation carrier; WT, wild type. ns, non-significance; *, *P*<0.05.

The phosphorylation level and expression of ERK, ECE1, EDN1 and cMyc were also detected in iPSCs-VSMCs. (**Figure 6d&e**) We observed that the inherent expression of pERK and EDN1 was high in the HPAH patient’s VSMCs, and progesterone could further induce the upregulation of EDN1, which was consistent with the previous results of PASMCs. Thus, we thought that EDN1 was the key protein in the pro-proliferative role of progesterone in BMPR2-mutated VSMCs. Although the inherent expression of cMyc and ECE1 was decreased in the HPAH patient’s VSMCs, it could be slightly upregulated by progesterone with no statistical difference.

### SM22-cre BMPR2 flox^+/-^ mice developed spontaneous PAH, which was further aggravated by progesterone

**Figure 7a** briefly showed the flowchart of drug intervention and phenotype detection of wild-type BMPR2 flox**^+/-^** female mice (WT mice) and SM22-cre BMPR2 flox^+/-^ female mice (CKO mice). Agarose gel electrophoresis demonstrated these two transgenic mice were successfully bred. (**Supplementary Figure 7a**) IHC results confirmed the decrease of BMPR2 in the pulmonary arteries of CKO mice. (**Supplementary Figure 7b&c**)

**Figure 7.**
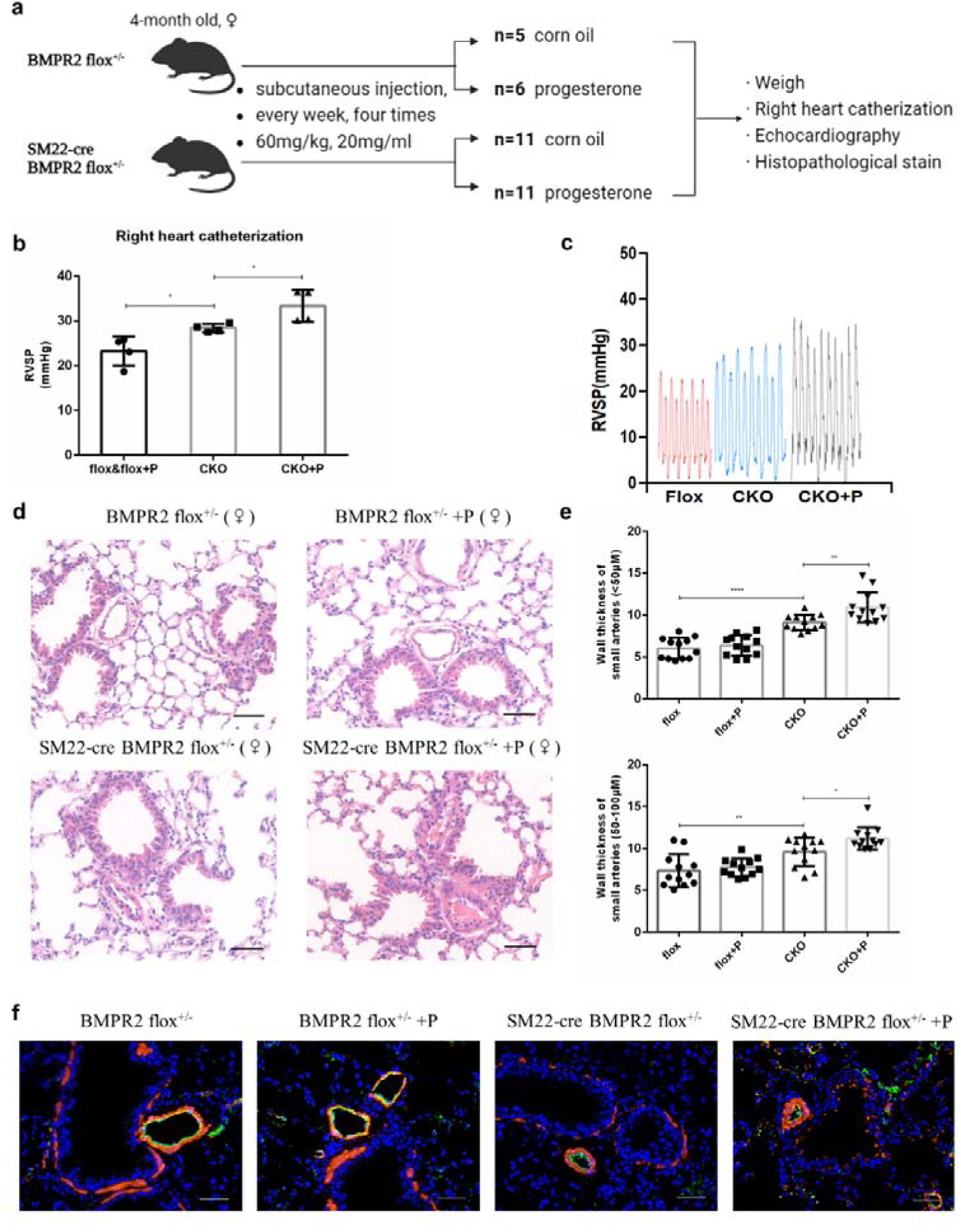
SM22-cre BMPR2 flox^+/-^ mice spontaneously developed PAH, which was further exacerbated by progesterone. (**a**) Time flowchart of animal intervention and phenotype detection. (**b-c**) Right heart catheterization experiments showed the increase of RVSP in CKO and CKO+P group. (n=4 per group) (**d-e**) H&E stain showed the vascular remodeling of pulmonary arterioles (<50μM and 50-100μM) in CKO and CKO+P group and the statistical analyses (4 mice per group, 3 fields per mouse). (**f**) IF stain of vWF (green) and αSMA (red) showed the proliferation and hypertrophy of PASMCs in CKO and CKO+P group. **Abbreviation:** ♀, female; RVSP, right ventricle systolic pressure; CKO, SM22-cre BMPR2 flox^+/-^ mice; P, progesterone; IF, immunofluorescence; vWF, von Willebrand factor; αSMA, smooth muscle actin. *, *P*<0.05; **, *P*<0.01; ****, *P*<0.0001.

The RHC finding proved that the RVSP of CKO mice was significantly increased compared with WT mice, and progesterone administration further up-regulated RVSP in CKO mice. (**Figure 7b–c**) H&E stain showed that progesterone aggravated the thickened wall of small and medium pulmonary arteries in CKO mice. (**Figure 7d–e**) Moreover, IF results manifested that pulmonary artery remodeling was mainly due to the proliferation and hypertrophy of PASMCs, rather than endothelium. (**Figure 7f**) However, no difference in weight was observed among these groups after one month of intervention of progesterone. (**Supplementary Figure 8a**) Besides, many results showed no development of RV remodeling in CKO group or CKO +progesterone group, including RV/LV+S, H&E stain and echocardiography. (**Supplementary Figure 8b-e**)

We further validated the related mechanisms of cellular experiments by Western blot in lung tissues. We found that pMYC, cMYC and EDN1 were elevated in CKO mice, and EDN1 was further up-regulated by progesterone in CKO mice. (**Figure 8a**) Similarly, IHC results showed a mild increase in the number of EDN1^+^ mesenchymal cells in the CKO group or CKO +progesterone groups, although low levels of EDN1 in pulmonary arteries. (**Figure 8b–c**)

**Figure 8.**
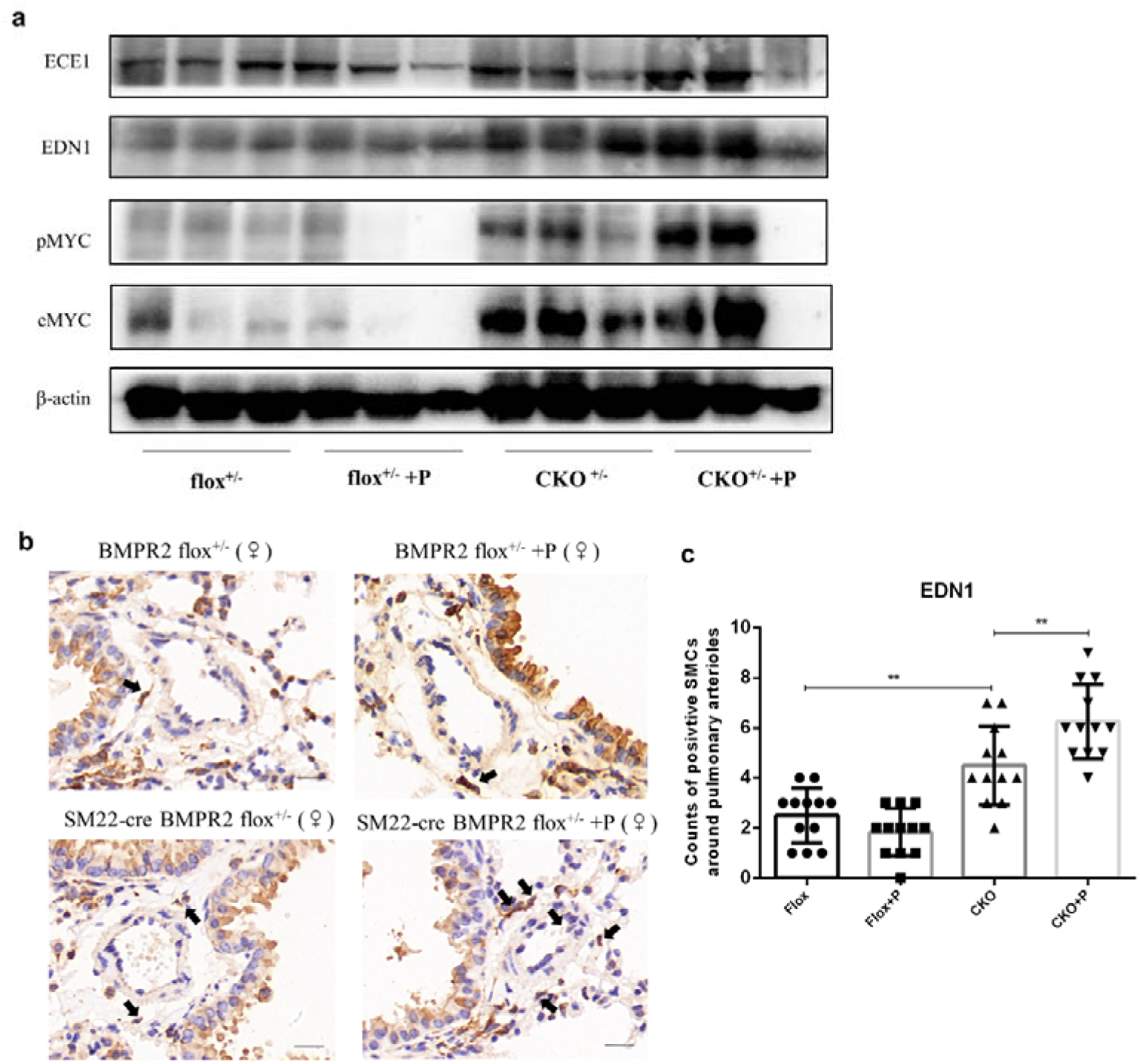
Validation of cellular mechanisms of phenotype changes and progesterone effects in SM22-cre BMPR2 flox^+/-^ mice. (**a**) Western blot of lung tissues of four groups (n=3 per group). (**b-c**) IHC stain of EDN1 and its statistical analysis of positive SMCs around pulmonary arterioles (3 mice per group, 4 fields per mouse). **, *P*<0.01

We also explored in CKO male mice and did not observe the remodeling of pulmonary arteries or RV in male at the age of 5 months, not like CKO female mice. (**Supplementary Figure 9a-b**) Furthermore, the upregulation of EDN1 and cMYC in lung tissues of CKO male mice compared to male WT mice was not as pronounced as females, suggesting the phenotypic difference and the underlying mechanism between different sexes of CKO mice. (**Supplementary Figure 9c**)

## Discussion

We innovatively demonstrated that progesterone promoted the proliferation of BMPR2-knockdown PASMCs by activating ERK pathway via PGR. Activated ERK not only upregulated the phosphorylation and elevation of cMYC, but also induced the transcription of EDN1 by promoting the nuclear entry of c-JUN and combination on the its promoter region. (**Supplementary Figure 10**) The above findings were further confirmed by using iPSCs-VSMCs. As for animal studies, we firstly reported that SM22-cre BMPR2 flox^+/-^ mice developed PAH spontaneously, which was further aggravated by exogenous progesterone.

Previous studies have identified some inducements of BMPR2-associated HPAH penetrance, including inflammation, somatic chromosome abnormalities, single nucleotide polymorphism of TGFβ1 and epigenetic alterations of BMPR2.^31–34^ As for the sex difference in penetrance, BMPR2-mutant female mice partly developed PAH spontaneously and had more severe PAH characteristics than males, and 16alpha-hydroxyoestrone (16OHE_1_) was determined as the crucial culprit.^35, 36^ However, no study focused on the effects of progesterone in this process, especially on BMPR2-deficient PASMCs and BMPR2-mutant transgenic mice. Our study fills this gap and goes further in explaining the high penetrance among women of reproductive age and its potential mechanisms.

*In vitro*, we demonstrated that progesterone had pro-proliferative effects on BMPR2-knockdown PASMCs but not on normal PASMCs. We believed this was due to the preservation of cellular self-regulation ability when the expression of BMPR2 was normal. We observed that the abundance of cMYC, ECE1, and EDN1 in PASMCs were significantly increased when BMPR2 declined, which puts PASMCs in an irritable state for proliferation. Progesterone stimulation further upregulated these proteins, thereby promoting the proliferation of PASMCs. Other studies also reported the irritable feature of BMPR2 mutation. For instance, BMPR2 heterozygosity decreased the expression of the antiapoptotic miR124-3p in PASMCs, which resulted in increased proliferation by accumulating unfolded proteins and enhancing PERK signaling under hypoxic conditions.^37^ Besides, BMPR2 mutation cause the drift of signaling pathways from smad1/5/8 axis to TGFβ-TAK1-MAPK axis under the stimulation of BMP4, thereby leading to proliferation via MAPK activation.^38, 39^

We revealed that progesterone activated the PGR-ERK axis, causing the proliferation of BMPR2-knockdown PASMCs. Other studies similarly showed that progesterone could activate it in human umbilical vein endothelial cells, which was reversed by the pretreatment with mifepristone, indicating that PGR was involved in this activation process.^40^ Regarding how progesterone activated the ERK pathway through PGR, a previous study in human breast cancer cells demonstrated that estrogen activated SRC/p21/ERK via estrogen receptor (ESR), and progesterone could induce the combination of the two domains of PGR with ESR, thereby activating the SRC/ERK pathway.^41^ Furthermore, our experiment was consistent with the previous results that activating the pERK-AP-1 axis could promote EDN1 transcription.^42, 43^ Advanced glycation end products significantly increased the binding activity of AP-1 in the promoter region of EDN1,^43^ and the mutation experiment indicated that the AP-1 binding site was an important cis-element for EDN1 expression induced by urea acids.^42^ As for iPSC-VSMCs experiments, several groups have proved that endothelial cells or cardiomyocytes derived from the BMPR2-mutated patient-specific iPSCs remained PAH-specific phenotypes and characteristics,^44, 45^ supporting the reliability of our results.

In line with our animal study, it has been reported that SM22-tet-BMPR2^delx4^ mice and SM22-rtTA x TetO7-BMPR2^R899X^ mice, in which BMPR2 was deleted in smooth muscle cells, spontaneously developed PAH after administration doxycycline. ^46, 47^ As the main downstream of BMPR2, SMAD1-deficiency in smooth muscle cells also predisposed mice to PAH at almost 10% chance.^48^ Importantly, we innovatively revealed that exogenous progesterone could aggravate the PAH phenotype of SM22-cre BMPR2 flox^+/-^ mice, indicating that progesterone might have extra influences different from estrogen.^35^ In CKO group and CKO +progesterone group, we observed the pulmonary vascular remodeling but not right heart hypertrophy, probably due to short observation time. Perhaps, we speculated that it might also be associated with the protective effects of progesterone on the heart.^49^

We suggested EDN1 was a key molecule for progesterone to promoted the proliferation of PASMCs in the absence of BMPR2, since it was significantly increased in primary PASMCs experiments, iPSC-VSMCs validation and transgenic animal studies. Similarly, previous literature reported that exogenous EDN1 could enhance the MAPK pathway and lead to cellular proliferation in not only normal PASMCs but also PASMCs from IPAH patients.^50, 51^ A study of mass spectrometry demonstrated that exogenous EDN1 enriched the pathways of proliferation, apoptosis and contraction in BMPR2-mutated PASMCs derived from PAH patients, compared with PASMCs from normal controls.^52^ In summary, the complex connection between BMPR2 and EDN1 might be related to the abnormal proliferation of PASMCs, synergistically leading to the occurrence and development of PAH. Our research firstly reported the regulation of EDN1 by sex hormones in the field of pulmonary vasculature.

Some limitations should be noticed in this study. First, it is not clear whether the source of commercially-purchased primary PASMCs in our cellular experiments was from male or female. Mair KM et al^53^ reported the sex-biased effect of 17β-estradiol on PASMCs, indicating that sex difference might affect the phenotypic sensitivity of progesterone on PASMCs. Second, Previous studies claimed that different 17β-estradiol metabolites had opposite effects on PASMCs, such as the pro-proliferative effect of 16-OHE_1_ and anti-proliferative effect of 2-methylestradiol (2ME).^54, 55^ Herein, we only studied the effects of progesterone without concerning its metabolites. Third, we used iPSCs from only 1 male patient with HPAH and 1 healthy person. In future, more iPSCs from HPAH patients carrying different BMPR2 mutation would be constructed to validate our results. Finally, we used CKO heterozygous mice, because we failed to breed CKO homozygous mice due to embryonic death. We did not choose the ERT2-SM22-Cre mouse, since tamoxifen as an ESR antagonist might interfere with the effects of progesterone.

## Conclusion

In BMPR2-deficient PASMCs, progesterone could promote the proliferation via PGR-ERK-cMYC/EDN1 axis. Exogenous progesterone could upregulate the lung expression of EDN1 and aggravate the phenotype of spontaneously-developed PAH in SM22-cre BMPR2 flox+/-mice. Progesterone might be an inducement of HPAH penetrance caused by BMPR2 mutation, accounting for the sex differential penetrance.

## Ethics approval and consent to participate

All mouse protocols and experimental procedures were approved by Shanghai Ruijin Hospital Institutional Animal Care and Use Committee. The establishment of iPS cell lines were approved by the Ethical Review Board of Shanghai Ruijin Hospital. (Approval number: (2019) BC001).

## Consent for publication

All the participants gave the informed consent for the collection of data and samples.

## Author contributions

Dong Liu and Wei-Ping Hu conceived and designed the study. Wei-Ping Hu and Si-Jin Zhang performed the experiments. Si-Min Xe and Li-Jun Fu participated in the experiment of iPSC construction. Wei-Ping Hu, Yong-Jie Ding and Jie Fang analyzed the data. Dong Liu and Shan-Qun Li funded this research. Wei-ping HU, Ling Zhou and Xiao Ge drafted the manuscript. Dong LIU, Jie-Ming QU, Qing-Yun Li, and Shan-Qun Li revised the draft of the manuscript. All authors read and approved the final manuscript for publication.

## Acknowledgement

We gave thanks to Dr. Bin-Feng He and Dr. Zong-Juan Li for their experimental guidance on ChIP assay. We also thanked Dr. Yu-Yi Liu and Prof. Jun Yang for the help in the culture and differentiation of iPSCs. We sincerely acknowledged Dr. Shu-Wen Qian and Prof. Qi-Qun Tang for generously providing the BMPR2 flox^+/-^ mice and the authentication method. Besides, we acknowledged Prof. Jian Wu and Prof. Jian-Guo Jia for their help in the animal experiments of RHC and echocardiography, respectively.

## Reference

1. Humbert M, Guignabert C, Bonnet S, Dorfmüller P, Klinger JR, Nicolls MR, Olschewski AJ, Pullamsetti SS, Schermuly RT, Stenmark KR, et al. Pathology and pathobiology of pulmonary hypertension: state of the art and research perspectives. Eur Respir J. 2019;53. doi: 10.1183/13993003.01887-2018

2. Kylhammar D, Hjalmarsson C, Hesselstrand R, Jansson K, Kavianipour M, Kjellstrom B, Nisell M, Soderberg S, Radegran G. Predicting mortality during long-term follow-up in pulmonary arterial hypertension. ERJ Open Res. 2021;7. doi: 10.1183/23120541.00837-2020

3. Machado RD, Southgate L, Eichstaedt CA, Aldred MA, Austin ED, Best DH, Chung WK, Benjamin N, Elliott CG, Eyries M, et al. Pulmonary Arterial Hypertension: A Current Perspective on Established and Emerging Molecular Genetic Defects. Hum Mutat. 2015;36:1113–1127. doi: 10.1002/humu.22904

4. Evans JD, Girerd B, Montani D, Wang XJ, Galie N, Austin ED, Elliott G, Asano K, Grunig E, Yan Y, et al. BMPR2 mutations and survival in pulmonary arterial hypertension: an individual participant data meta-analysis. Lancet Respir Med. 2016;4:129–137. doi: 10.1016/S2213-2600(15)00544-5

5. Andruska A, Spiekerkoetter E. Consequences of BMPR2 Deficiency in the Pulmonary Vasculature and Beyond: Contributions to Pulmonary Arterial Hypertension. Int J Mol Sci. 2018;19. doi: 10.3390/ijms19092499

6. Larkin EK, Newman JH, Austin ED, Hemnes AR, Wheeler L, Robbins IM, West JD, Phillips JA, 3rd, Hamid R, Loyd JE. Longitudinal analysis casts doubt on the presence of genetic anticipation in heritable pulmonary arterial hypertension. Am J Respir Crit Care Med. 2012;186:892–896. doi: 10.1164/rccm.201205-0886OC

7. McGoon MD, Benza RL, Escribano-Subias P, Jiang X, Miller DP, Peacock AJ, Pepke-Zaba J, Pulido T, Rich S, Rosenkranz S, et al. Pulmonary arterial hypertension: epidemiology and registries. J Am Coll Cardiol. 2013;62:D51–59. doi: 10.1016/j.jacc.2013.10.023

8. Santos-Ferreira C, Cardoso D, Paiva B, Baptista R. Pulmonary arterial hypertension unveils itself: a cancer-like progression – a case report. Eur Heart J Case Rep. 2021;5:ytab149. doi: 10.1093/ehjcr/ytab149

9. Elkus R, Popovich J, Jr. Respiratory physiology in pregnancy. Clin Chest Med. 1992;13:555–565.

10. Chaudhary KR, Deng Y, Yang A, Cober ND, Stewart DJ. Penetrance of Severe Pulmonary Arterial Hypertension in Response to Vascular Endothelial Growth Factor Receptor 2 Blockade in a Genetically Prone Rat Model Is Reduced by Female Sex. J Am Heart Assoc. 2021;10:e019488. doi: 10.1161/JAHA.120.019488

11. Tofovic PS, Zhang X, Petrusevska G. Progesterone inhibits vascular remodeling and attenuates monocrotaline-induced pulmonary hypertension in estrogen-deficient rats. Prilozi. 2009;30:25–44.

12. Zhang YX, Wang L, Lu WZ, Yuan P, Wu WH, Zhou YP, Zhao QH, Zhang SJ, Li Y, Wu T, et al. Association Between High FSH, Low Progesterone, and Idiopathic Pulmonary Arterial Hypertension in Women of Reproductive Age. Am J Hypertens. 2020;33:99–105. doi: 10.1093/ajh/hpz143

13. Tabula Muris C, Overall c, Logistical c, Organ c, processing, Library p, sequencing, Computational data a, Cell type a, Writing g, et al. Single-cell transcriptomics of 20 mouse organs creates a Tabula Muris. Nature. 2018;562:367–372. doi: 10.1038/s41586-018-0590-4

14. Kiprono LV, Wallace K, Moseley J, Martin J, Jr., Lamarca B. Progesterone blunts vascular endothelial cell secretion of endothelin-1 in response to placental ischemia. Am J Obstet Gynecol. 2013;209:44 e41–46. doi: 10.1016/j.ajog.2013.03.032

15. Ueda K, Lu Q, Baur W, Aronovitz MJ, Karas RH. Rapid estrogen receptor signaling mediates estrogen-induced inhibition of vascular smooth muscle cell proliferation. Arterioscler Thromb Vasc Biol. 2013;33:1837–1843. doi: 10.1161/ATVBAHA.112.300752

16. Deng L, Blanco FJ, Stevens H, Lu R, Caudrillier A, McBride M, McClure JD, Grant J, Thomas M, Frid M, et al. MicroRNA-143 Activation Regulates Smooth Muscle and Endothelial Cell Crosstalk in Pulmonary Arterial Hypertension. Circ Res. 2015;117:870–883. doi: 10.1161/CIRCRESAHA.115.306806

17. Hu WP, Xie L, Hao SY, Wu QH, Xiang GL, Li SQ, Liu D. Protective effects of progesterone on pulmonary artery smooth muscle cells stimulated with Interleukin 6 via blocking the shuttling and transcriptional function of STAT3. Int Immunopharmacol. 2022;102:108379. doi: 10.1016/j.intimp.2021.108379

18. Zhang SJ, Lian TY, Zhu XJ, Lu D, Xu XQ, Jiang X, Jing ZC. Derivation of an induced pluripotent stem cell line (PUMCHi003-A) from a patient with pulmonary arterial hypertension carrying heterozygous mutation in PTGIS gene. Stem Cell Res. 2020;46:101875. doi: 10.1016/j.scr.2020.101875

19. Patsch C, Challet-Meylan L, Thoma EC, Urich E, Heckel T, O’Sullivan JF, Grainger SJ, Kapp FG, Sun L, Christensen K, et al. Generation of vascular endothelial and smooth muscle cells from human pluripotent stem cells. Nat Cell Biol. 2015;17:994–1003. doi: 10.1038/ncb3205

20. Li Z, Luo G, Hu WP, Hua JL, Geng S, Chu PK, Zhang J, Wang H, Yu XF. Mediated Drug Release from Nanovehicles by Black Phosphorus Quantum Dots for Efficient Therapy of Chronic Obstructive Pulmonary Disease. Angew Chem Int Ed Engl. 2020;59:20568–20576. doi: 10.1002/anie.202008379

21. Qian S, Pan J, Su Y, Tang Y, Wang Y, Zou Y, Zhao Y, Ma H, Zhang Y, Liu Y, et al. BMPR2 promotes fatty acid oxidation and protects white adipocytes from cell death in mice. Commun Biol. 2020;3:200. doi: 10.1038/s42003-020-0928-y

22. Kim O, Park EY, Kwon SY, Shin S, Emerson RE, Shin YH, DeMayo FJ, Lydon JP, Coffey DM, Hawkins SM, et al. Targeting progesterone signaling prevents metastatic ovarian cancer. Proc Natl Acad Sci U S A. 2020;117:31993–32004. doi: 10.1073/pnas.2013595117

23. Jiang T, Hu W, Zhang S, Ren C, Lin S, Zhou Z, Wu H, Yin J, Tan L. Fibroblast growth factor 10 attenuates chronic obstructive pulmonary disease by protecting against glycocalyx impairment and endothelial apoptosis. Respir Res. 2022;23:269. doi: 10.1186/s12931-022-02193-5

24. Asati V, Mahapatra DK, Bharti SK. PI3K/Akt/mTOR and Ras/Raf/MEK/ERK signaling pathways inhibitors as anticancer agents: Structural and pharmacological perspectives. Eur J Med Chem. 2016;109:314–341. doi: 10.1016/j.ejmech.2016.01.012

25. Dhanasekaran R, Deutzmann A, Mahauad-Fernandez WD, Hansen AS, Gouw AM, Felsher DW. The MYC oncogene – the grand orchestrator of cancer growth and immune evasion. Nat Rev Clin Oncol. 2022;19:23–36. doi: 10.1038/s41571-021-00549-2

26. Galie N, Channick RN, Frantz RP, Grunig E, Jing ZC, Moiseeva O, Preston IR, Pulido T, Safdar Z, Tamura Y, et al. Risk stratification and medical therapy of pulmonary arterial hypertension. Eur Respir J. 2019;53. doi: 10.1183/13993003.01889-2018

27. Lee SH, Hu LL, Gonzalez-Navajas J, Seo GS, Shen C, Brick J, Herdman S, Varki N, Corr M, Lee J, et al. ERK activation drives intestinal tumorigenesis in Apc(min/+) mice. Nat Med. 2010;16:665–670. doi: 10.1038/nm.2143

28. Han J, Zhang Y, Xu J, Zhang T, Wang H, Wang Z, Jiang Y, Zhou L, Yang M, Hua Y, et al. Her4 promotes cancer metabolic reprogramming via the c-Myc-dependent signaling axis. Cancer Lett. 2021;496:57–71. doi: 10.1016/j.canlet.2020.10.008

29. Xu D, Emoto N, Giaid A, Slaughter C, Kaw S, deWit D, Yanagisawa M. ECE-1: a membrane-bound metalloprotease that catalyzes the proteolytic activation of big endothelin-1. Cell. 1994;78:473–485. doi: 10.1016/0092-8674(94)90425-1

30. Martinez-Miguel P, Medrano-Andres D, Griera-Merino M, Ortiz A, Rodriguez-Puyol M, Rodriguez-Puyol D, Lopez-Ongil S. Tweak up-regulates endothelin-1 system in mouse and human endothelial cells. Cardiovasc Res. 2017;113:207–221. doi: 10.1093/cvr/cvw239

31. Liu D, Yan Y, Chen JW, Yuan P, Wang XJ, Jiang R, Wang L, Zhao QH, Wu WH, Simonneau G, et al. Hypermethylation of BMPR2 Promoter Occurs in Patients with Heritable Pulmonary Arterial Hypertension and Inhibits BMPR2 Expression. Am J Respir Crit Care Med. 2017;196:925–928. doi: 10.1164/rccm.201611-2273LE

32. Phillips JA, 3rd, Poling JS, Phillips CA, Stanton KC, Austin ED, Cogan JD, Wheeler L, Yu C, Newman JH, Dietz HC, et al. Synergistic heterozygosity for TGFbeta1 SNPs and BMPR2 mutations modulates the age at diagnosis and penetrance of familial pulmonary arterial hypertension. Genet Med. 2008;10:359–365. doi: 10.1097/GIM.0b013e318172dcdf

33. Aldred MA, Comhair SA, Varella-Garcia M, Asosingh K, Xu W, Noon GP, Thistlethwaite PA, Tuder RM, Erzurum SC, Geraci MW, et al. Somatic chromosome abnormalities in the lungs of patients with pulmonary arterial hypertension. Am J Respir Crit Care Med. 2010;182:1153–1160. doi: 10.1164/rccm.201003-0491OC

34. Sawada H, Saito T, Nickel NP, Alastalo TP, Glotzbach JP, Chan R, Haghighat L, Fuchs G, Januszyk M, Cao A, et al. Reduced BMPR2 expression induces GM-CSF translation and macrophage recruitment in humans and mice to exacerbate pulmonary hypertension. J Exp Med. 2014;211:263–280. doi: 10.1084/jem.20111741

35. Fessel JP, Chen X, Frump A, Gladson S, Blackwell T, Kang C, Johnson J, Loyd JE, Hemnes A, Austin E, et al. Interaction between bone morphogenetic protein receptor type 2 and estrogenic compounds in pulmonary arterial hypertension. Pulm Circ. 2013;3:564–577. doi: 10.1086/674312

36. Mair KM, Harvey KY, Henry AD, Hillyard DZ, Nilsen M, MacLean MR. Obesity alters oestrogen metabolism and contributes to pulmonary arterial hypertension. Eur Respir J. 2019;53. doi: 10.1183/13993003.01524-2018

37. Shimizu T, Higashijima Y, Kanki Y, Nakaki R, Kawamura T, Urade Y, Wada Y. PERK inhibition attenuates vascular remodeling in pulmonary arterial hypertension caused by BMPR2 mutation. Sci Signal. 2021;14. doi: 10.1126/scisignal.abb3616

38. Nasim MT, Ogo T, Chowdhury HM, Zhao L, Chen CN, Rhodes C, Trembath RC. BMPR-II deficiency elicits pro-proliferative and anti-apoptotic responses through the activation of TGFbeta-TAK1-MAPK pathways in PAH. Hum Mol Genet. 2012;21:2548–2558. doi: 10.1093/hmg/dds073

39. Dewachter L, Adnot S, Guignabert C, Tu L, Marcos E, Fadel E, Humbert M, Dartevelle P, Simonneau G, Naeije R, et al. Bone morphogenetic protein signalling in heritable versus idiopathic pulmonary hypertension. Eur Respir J. 2009;34:1100–1110. doi: 10.1183/09031936.00183008

40. Hsu SP, Lee WS. Progesterone receptor activation of extranuclear signaling pathways in regulating p53 expression in vascular endothelial cells. Mol Endocrinol. 2011;25:421–432. doi: 10.1210/me.2010-0424

41. Vicent GP, Ballare C, Zaurin R, Saragueta P, Beato M. Chromatin remodeling and control of cell proliferation by progestins via cross talk of progesterone receptor with the estrogen receptors and kinase signaling pathways. Ann N Y Acad Sci. 2006;1089:59–72. doi: 10.1196/annals.1386.025

42. Chao HH, Liu JC, Lin JW, Chen CH, Wu CH, Cheng TH. Uric acid stimulates endothelin-1 gene expression associated with NADPH oxidase in human aortic smooth muscle cells. Acta Pharmacol Sin. 2008;29:1301–1312. doi: 10.1111/j.1745-7254.2008.00877.x

43. Adamopoulos C, Piperi C, Gargalionis AN, Dalagiorgou G, Spilioti E, Korkolopoulou P, Diamanti-Kandarakis E, Papavassiliou AG. Advanced glycation end products upregulate lysyl oxidase and endothelin-1 in human aortic endothelial cells via parallel activation of ERK1/2-NF-kappaB and JNK-AP-1 signaling pathways. Cell Mol Life Sci. 2016;73:1685–1698. doi: 10.1007/s00018-015-2091-z

44. Gu M, Shao NY, Sa S, Li D, Termglinchan V, Ameen M, Karakikes I, Sosa G, Grubert F, Lee J, et al. Patient-Specific iPSC-Derived Endothelial Cells Uncover Pathways that Protect against Pulmonary Hypertension in BMPR2 Mutation Carriers. Cell Stem Cell. 2017;20:490–504 e495. doi: 10.1016/j.stem.2016.08.019

45. Du M, Jiang H, Liu H, Zhao X, Zhou Y, Zhou F, Piao C, Xu G, Ma F, Wang J, et al. Single-cell RNA sequencing reveals that BMPR2 mutation regulates right ventricular function via ID genes. Eur Respir J. 2021. doi: 10.1183/13993003.00327-2021

46. West J, Harral J, Lane K, Deng Y, Ickes B, Crona D, Albu S, Stewart D, Fagan K. Mice expressing BMPR2R899X transgene in smooth muscle develop pulmonary vascular lesions. Am J Physiol Lung Cell Mol Physiol. 2008;295:L744–755. doi: 10.1152/ajplung.90255.2008

47. West J, Fagan K, Steudel W, Fouty B, Lane K, Harral J, Hoedt-Miller M, Tada Y, Ozimek J, Tuder R, et al. Pulmonary hypertension in transgenic mice expressing a dominant-negative BMPRII gene in smooth muscle. Circ Res. 2004;94:1109–1114. doi: 10.1161/01.RES.0000126047.82846.20

48. Han C, Hong KH, Kim YH, Kim MJ, Song C, Kim MJ, Kim SJ, Raizada MK, Oh SP. SMAD1 deficiency in either endothelial or smooth muscle cells can predispose mice to pulmonary hypertension. Hypertension. 2013;61:1044–1052. doi: 10.1161/HYPERTENSIONAHA.111.199158

49. Frump AL, Albrecht M, Yakubov B, Breuils-Bonnet S, Nadeau V, Tremblay E, Potus F, Omura J, Cook T, Fisher A, et al. 17beta-Estradiol and estrogen receptor alpha protect right ventricular function in pulmonary hypertension via BMPR2 and apelin. J Clin Invest. 2021;131. doi: 10.1172/JCI129433

50. Maruyama H, Dewachter C, Belhaj A, Rondelet B, Sakai S, Remmelink M, Vachiery JL, Naeije R, Dewachter L. Endothelin-Bone morphogenetic protein type 2 receptor interaction induces pulmonary artery smooth muscle cell hyperplasia in pulmonary arterial hypertension. J Heart Lung Transplant. 2015;34:468–478. doi: 10.1016/j.healun.2014.09.011

51. Yamanaka R, Otsuka F, Nakamura K, Yamashita M, Otani H, Takeda M, Matsumoto Y, Kusano KF, Ito H, Makino H. Involvement of the bone morphogenetic protein system in endothelin-and aldosterone-induced cell proliferation of pulmonary arterial smooth muscle cells isolated from human patients with pulmonary arterial hypertension. Hypertens Res. 2010;33:435–445. doi: 10.1038/hr.2010.16

52. Yao C, Yu J, Taylor L, Polgar P, McComb ME, Costello CE. Protein Expression by Human Pulmonary Artery Smooth Muscle Cells Containing a BMPR2 Mutation and the Action of ET-1 as Determined by Proteomic Mass Spectrometry. Int J Mass Spectrom. 2015;378:347–359. doi: 10.1016/j.ijms.2014.10.006

53. Mair KM, Yang XD, Long L, White K, Wallace E, Ewart MA, Docherty CK, Morrell NW, MacLean MR. Sex affects bone morphogenetic protein type II receptor signaling in pulmonary artery smooth muscle cells. Am J Respir Crit Care Med. 2015;191:693–703. doi: 10.1164/rccm.201410-1802OC

54. White K, Johansen AK, Nilsen M, Ciuclan L, Wallace E, Paton L, Campbell A, Morecroft I, Loughlin L, McClure JD, et al. Activity of the estrogen-metabolizing enzyme cytochrome P450 1B1 influences the development of pulmonary arterial hypertension. Circulation. 2012;126:1087–1098. doi: 10.1161/CIRCULATIONAHA.111.062927

55. Hao S, Jiang L, Fu C, Wu X, Liu Z, Song J, Lu H, Wu X, Li S. 2-Methoxyestradiol attenuates chronic-intermittent-hypoxia-induced pulmonary hypertension through regulating microRNA-223. J Cell Physiol. 2019;234:6324–6335. doi: 10.1002/jcp.27363

